# Joint analysis of area and thickness as a replacement for the analysis of cortical volume

**DOI:** 10.1101/074666

**Authors:** Anderson M. Winkler, Douglas N. Greve, Knut J. Bjuland, Thomas E. Nichols, Mert R. Sabuncu, Asta K. Håberg, Jon Skranes, Lars M. Rimol

## Abstract

Cortical surface area is an increasingly used brain morphology metric that is ontogenetically and phylogenetically distinct from cortical thickness and offers a separate index of neurodevelopment and disease. However, the various existing methods for assessment of cortical surface area from magnetic resonance images have never been systematically compared. We show that the surface area method implemented in FreeSurfer corresponds closely to the exact, but computationally more demanding, mass-conservative (pyc-nophylactic) method, provided that images are smoothed. Thus, the data produced by this method can be interpreted as estimates of cortical surface area, as opposed to areal expansion. In addition, focusing on the joint analysis of thickness and area, we compare an improved, analytic method for measuring cortical volume to a permutation based non-parametric combination (NPC) method. We use the methods to analyse area, thickness and volume in young adults born preterm with very low birth weight, and show that NPC analysis is a more sensitive option for studying joint effects on area and thickness, giving equal weight to variation in both of these two morphological features.

## 1. Introduction

It has been suggested that biological processes that drive horizontal (tangential) and vertical (radial) development of the cerebral cortex are separate from each other (Rakic, 1988; Geschwind and Rakic, 2013), influencing cortical area and thickness independently. These two indices of cerebral morphology are uncorrelated genetically (Panizzon et al., 2009; Winkler et al., 2010), are each influenced by regionally distinct genetic factors (Schmitt et al., 2008; Rimol et al., 2010b; Chen et al., 2012, 2015), follow different trajectories over the lifespan (O’Leary et al., 2007; Hogstrom et al., 2013; Fjell et al., 2015), and are differentially associated with cognitive abilities and disorders (Schnack et al., 2015; Noble et al., 2015; Lee et al., 2016; Vuoksimaa et al., 2016). Moreover, it is cortical area, not thickness, that differs substantially across species (Rakic, 1995). These findings give prominence to the use of surface area in studies of cortical morphology and its relationship to function. However, a number of approaches and terminologies exist for its assessment, which have not been studied in detail or compared directly, making interpretation and comparison between studies challenging. A first objective of this paper is to compare the methods for the analysis of cortical area, in particular interpolation between surfaces at different resolutions, and to provide recommendations for users.

A second objective of the paper is to demonstrate that a statistical joint analysis of cortical thickness and surface area, using the recently proposed Non-Parametric Combination (NPC; Pesarin and Salmaso, 2010b; Winkler et al., 2016b), can be used to investigate factors affecting cortical morphology. While analyzing cortical thickness and cortical area separately improves specificity over combined metrics such as cortical volume (Rimol et al., 2012), it may still be of interest to jointly analyze these two measurements so as to increase power to detect effects on thickness and area simultaneously. In principle, this could be accomplished through the analysis of cortical volume, which commingles thickness and area. Indeed, volume is a popular metric, thanks mainly to the wide use of voxel-based morphometry (VBM; Ashburner and Friston, 2000; Good et al., 2001; Douaud et al., 2007), despite a series of well documented disadvantages (Davatzikos, 2004; Ashburner, 2009). In surface-based approaches, cortical volume is measured as the product of cortical thickness and surface area at each location across the cortical mantle. However, here we demonstrate that this multiplicative method incurs severe bias, the direction of which varies according to the local geometry of the cortex. In order to conduct a fair comparison of surface-based cortical volume analysis and joint analysis with NPC, we propose a novel, geometrically exact, analytic solution to the measurement of cortical volume, which does not suffer from such bias, and compare this improved cortical volume method to analysis with NPC.

### 1.1. Cortical surface area

Using continuous (vertexwise) cortical maps to compare surface area across subjects has the advantage that, unlike approaches based on regions of interest (ROI), it does not depend on the effects of interest mapping onto a previously defined ROI scheme. However, few studies using continuous maps have offered insight into the procedures adopted. Sometimes the methods were described in terms of areal expansion/contraction, as opposed to surface area itself, and different definitions of areal expansion/contraction have been used, e.g., relative to the contra-lateral hemisphere (Lyttelton et al., 2009), to some earlier point in time (Hill et al., 2010), to a control group (Palaniyappan et al., 2011), or in relation to a standard brain, possibly the default brain (average or atlas) used in the respective software package (Joyner et al., 2009; Rimol et al., 2010a, 2012; Chen et al., 2011, 2012; Vuoksimaa et al., 2016); other studies considered linear distances as proxies for expansion/contraction (Sun et al., 2009a,b). Some of the studies that used a default brain as reference used nearest neighbour interpolation followed by smoothing, which, as we show below, assesses cortical area itself, but described the measurements in terms of areal expansion (Joyner et al., 2009; Rimol et al., 2010a, 2012).

Surface area analyses depend on registration of the cortical surface and interpolation to a common resolution; such resampling must preserve the amount of area at local, regional and global scales, i.e., it must be mass-conservative. This means that the choice of interpolation method is crucial. A well-known interpolation method is *nearest-neighbour*, which can be enhanced by correction for stretches and shrinkages of the surface during registration, as available in the function `mris_preproc`, part of the FreeSurfer (FS) software package.^1^ Another approach is *retessellation* of the mesh of the individual subject to the geometry of a common grid, as proposed by Saad et al. (2004) as a way to produce meshes with similar geometry across subjects. Even though the method has been mostly used to compute areal expansion, it can be used for surface area itself, as well as for other areal quantities. A third approach is the use of the barycentric coordinates of each vertex with reference to the vertices of the common grid to *redistribute* the areal quantities, in an approximately mass conservative process. Lastly, a strategy for analysis of areal quantities using a *pycnophylactic* (mass-preserving) interpolation method, which addresses the above concerns, but that is computationally intensive, was presented in Winkler et al. (2012) (Table 1).

**Table 1:** Overview of the four different methods to interpolate surface area and areal quantities. A detailed description is in the Materials and Methods section.

| **Method** | **Description** |
| --- | --- |
| Nearest-neighbour | Nearest-neighbour interpolation of areal quantities on the sphere, followed by Jacobian correction. |
| Retessellation | Barycentric interpolation on the sphere of the native vertex coordinates. |
| Redistributive | Vertexwise redistribution of areal quantities based on barycentric coordinates of the source in relation to the target. |
| Pycnophylactic | Mass-conservative facewise interpolation method that uses the overlapping areas between faces of source and target. |

### 1.2. Measuring volume and other areal quantities

The volume of cortical grey matter is also an areal quantity, which therefore requires mass-conservative interpolation methods. Volume can be estimated through the use of voxelwise partial volume effects using a volume-based representation of the brain, such as in VBM, or from a surface representation, in which it can be measured as the amount of tissue present between the surface placed at the site of the pia mater, and the surface at the interface between gray and white matter. If the area of either of these surfaces is known, or if the area of a mid-surface, i.e., the surface running half-distance between pial and white surfaces (van Essen, 2005) is known, an estimate of the volume can be obtained by multiplying, at each vertex, area by thickness. This procedure, while providing a reasonable approximation that improves over voxel-based measurements for being less susceptible to various artefacts (for a discussion of artefacts in VBM, see Ashburner, 2009), is still problematic as it underestimates the volume of tissue that is external to the convexity of the surface, and overestimates volume that is internal to it; both cases are undesirable, and cannot be solved by merely resorting to using an intermediate surface as the mid-surface (Figure 1a). Here a different approach is proposed: each face of the white surface and its matching face in the pial surface are used to define an oblique truncated pyramid, the volume of which is computed analytically, without introducing additional error other than what is intrinsic to the placement and resolution of these surfaces (Figure 1b for a 2-D schema and Figure 2 for a similar in 3-D).

**Figure 1:**
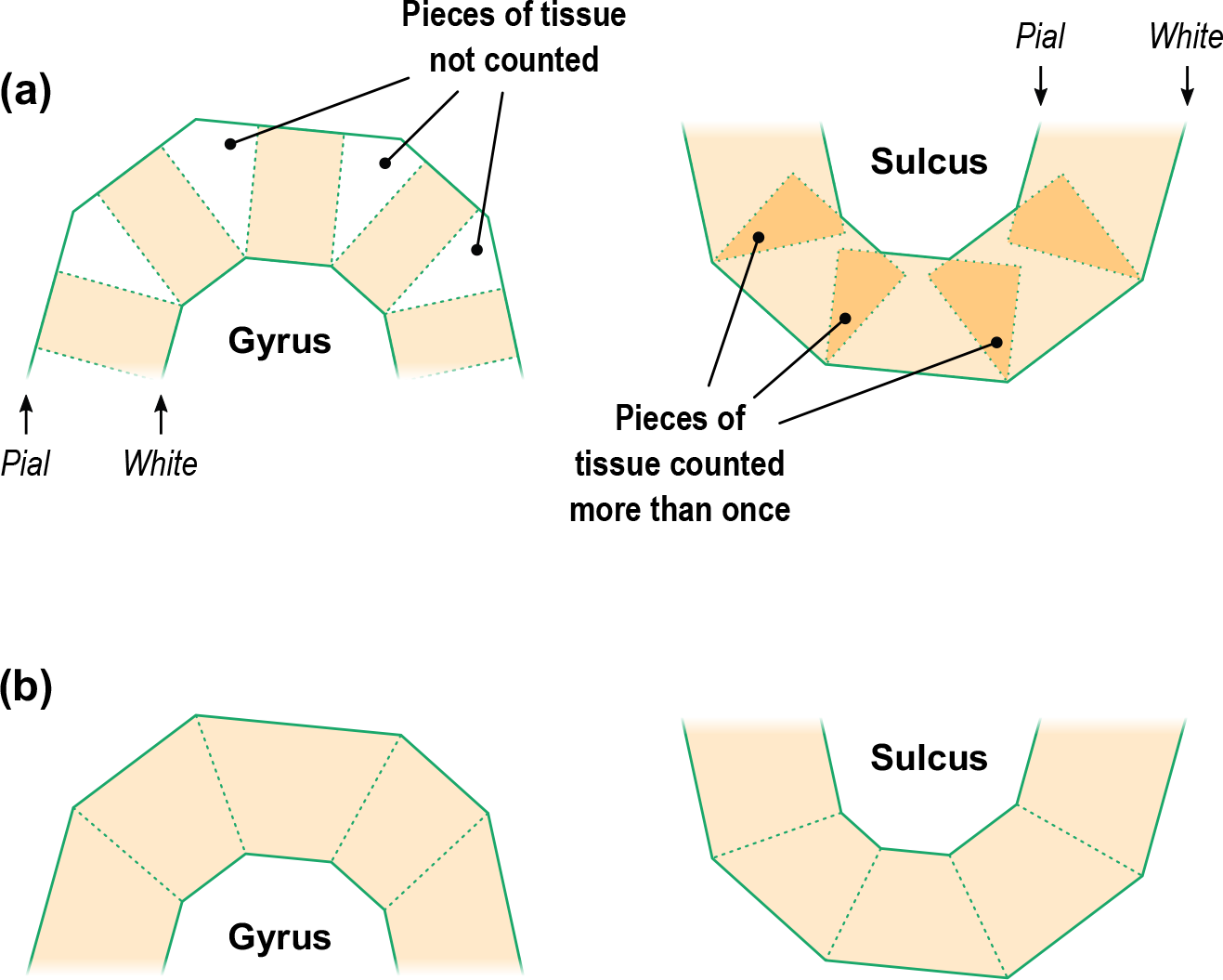
A diagram in two dimensions of the problem of measuring the cortical volume. (*a*) If volume is computed using multiplication of thickness by area, considerable amount of tissue is left unmeasured in the gyri, or measured repeatedly in sulci. The problem is minimised, but not solved, with the use of the mid-surface. (*b*) Instead, vertex coordinates can be used to compute analytically the volume of tissue between matching faces of white and pial surfaces, leaving no tissue under- or over-represented.

**Figure 2:**
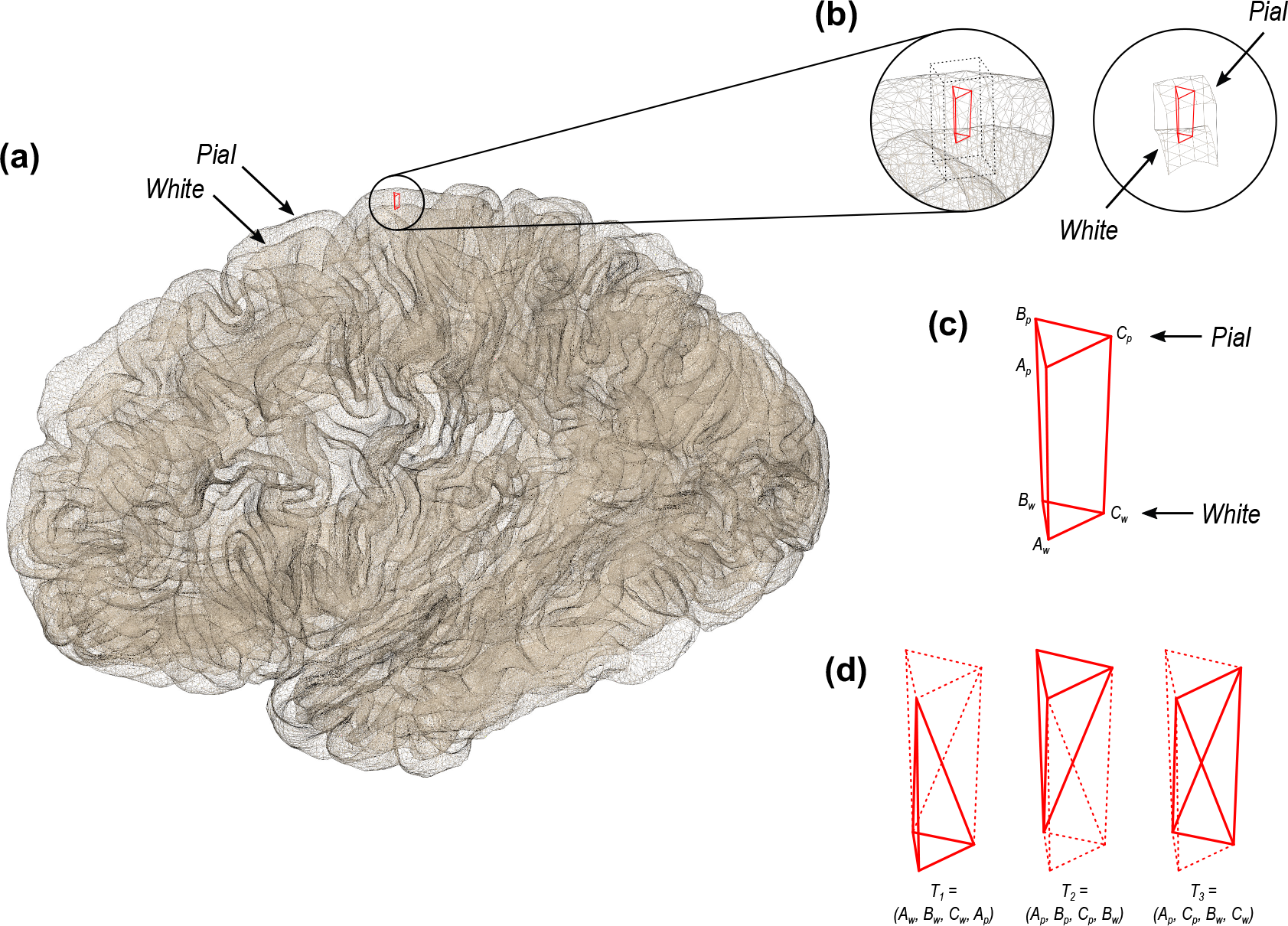
(*a*) In the surface representation, the cortex is limited internally by the white and externally by the pial surface. (*b*) and (*c*) These two surfaces have matching vertices that can be used to delineate an oblique truncated triangular pyramid. (*d*) The six vertices of this pyramid can be used to define three tetrahedra, the volumes of which are computed analytically.

Quantitative measurements, such as from positron emission tomography (PET), cerebral blood flow, cerebral blood volume, the mass, or number of molecules of a given compound (Leahy and Qi, 2000; van den Hoff, 2005), are all areal quantities whenever these are expressed in absolute quantities. Likewise, cerebral blood flow and volume obtained using methods based on magnetic resonance imaging (MRI), such as arterial spin labelling (ASL), as well as other forms of quantitative MRI, as those involving contrast enhancement (Parker and Padhani, 2003), quantitative magnetisation transfer (Levesque et al., 2010; Harrison et al., 2015), or quantitative assessment of myelination, are also areal quantities that require mass conservation when measured in absolute terms. The methods used for statistical analysis surface area can be applied for these areal quantities as well.

### 1.3. Non-parametric combination (NPC)

We argue that analysing thickness and area jointly offers important advantages over using cortical volume, regardless of how the latter is measured. The permutation-based Non-Parametric Combination (NPC; Pesarin and Salmaso, 2010b; Winkler et al., 2016b) supplies a test for directional as well as two-tailed hypotheses, which has been proven to be more powerful than classical multivariate tests (Pesarin and Salmaso, 2010a). The NPC consists of, in a first phase, testing separately hypotheses on each available metric (here thickness and area) using permutations that are performed in synchrony; these tests are termed *partial tests.* The resulting statistics for every permutation are recorded, allowing an estimate of the complete empirical cumulative distribution function (cdf) to be constructed for each metric. In a second phase, the empirical p-values are combined, for each permutation, into a *joint statistic.* As the joint statistic is produced from the previous permutations that have been recorded, an estimate of its empirical cdf is immediately known, and therefore its p-value. The test is based on minimal assumptions, mainly that of exchangeability, that is, swapping one datum for another keeps the data just as likely. Independence among metrics or partial tests is not assumed by NPC: the synchronised permutations implicitly capture eventual dependencies. This is particularly important when investigating cortical area and thickness, since shared environmental effects may affect area and thickness simultaneously.

## 2. Method

We apply the methods to a cohort of adults born preterm with very low birth weight (⩽ 1500g, VLBW), and a set of coetaneous controls born at the same hospital and period. At the age 20, from an initial group of 121 VLBW and 122 control subjects, a total of 41 VLBW and 59 controls consented to participate and had usable MRI data. Details about the sample can be found in Martinussen et al. (2005); Skranes et al. (2007). The local Regional Committee for Medical Research Ethics approved the study protocol (Norwegian Health Region IV; REK project number: 4.2005.2605). Previous studies of the preterm population have shown reduced cortical surface area in multiple frontal, temporal and parieto-occipital regions, as well as increased frontal lobe and reduced parietal lobe cortical thickness. This combination of reductions and increases, in partly overlapping regions, makes the present sample well-suited for a demonstration of a joint analysis of surface area and cortical thickness. Using this data we evaluate the four different interpolation methods (nearest neighbour, retessellation, redistributive and pycnophylactic) differ, how they interact with different resolutions (shown in the Supplementary Material only), the two ways of measuring volume (the product method and the analytic method), and finally, we demonstrate benefits of NPC over cortical volume for the investigation of differences in cortical morphology between the two groups. We note that comparisons among interpolation methods depend only on algorithmic and geometric differences between them, not interacting with particular features of this or any sample, such that the results are generalisable, and not affected by pathology.

### 2.1. Data acquisition and surface reconstruction

MRI scanning was performed on a 1.5 T Siemens MAGNETOM Symphony scanner. Two sagittal *T*_1_-weighted magnetization prepared rapid gradient echo (MPRAGE) scans were acquired (TE/TR/TI = 3.45/2730/1000 ms, flip angle = 7°, voxel size = 1 × 1 × 1.33 mm). Surfaces were reconstructed using the FreeSurfer software package (version 5.3.0; Dale et al., 1999; Fischl et al., 1999a), and an overview of the whole process is in Supplementary Material §1. We used the FreeSurfer software suite but similar methods for surface reconstruction exist in other software packages (Mangin et al., 1995; van Essen et al., 2001; Kim et al., 2005), and the present comparisons of interpolation methods and methods of volume measurement, as well as analysis with NPC, are not specific to FreeSurfer.

### 2.2. Measurement of areal quantities

Areal quantities are measured in native space, i.e., before registration. For the retessellation method, the measurement is made in native space after the surface has been reconstructed to a common grid; nearest neighbour, redistributive, and pycnophylactic use native space measurements with the original, subject-specific mesh geometry.

#### Cortical area

For a triangular face *ABC* of the surface representation, with vertex coordinates **a** = [*x*_*A*_ *y*_*A*_ *z*_*A*_]′, **b** = [*x*_*B*_ *y*_*B*_ *z*_*B*_]′, and **c** = [*x*_*C*_ *y*_*C*_ *z*_*C*_]′, the area is |**u** × **v**|/2, where **u** = **a** − **c**, **v** = **b** − **c**, × represents the cross product, and the bars | | represent the vector norm. Although the area per face (i.e., the *facewise* area) can be used in subsequent steps, it remains the case that most software packages can only deal with values assigned to each vertex of the mesh (i.e., *vertexwise*). Conversion from facewise to vertexwise is achieved by assigning to each vertex one-third of the sum of the areas of all faces that have that vertex in common (Winkler et al., 2012).

#### Cortical volume

The conventional method for computing surface-based volume consists of computing the area at each vertex as above, then multiplying this value by the thickness at that vertex, in a procedure that leaves tissue under- or over-represented in gyri and sulci (Figure 1). We propose that, instead, volumes are computed using the three vertices that define a face in the white surface and the three matching vertices in the pial surface, defining an *oblique truncated triangular pyramid*, which in its turn is subdivided into three tetrahedra. The volumes of these are computed analytically, summed, and assigned to each face of the surface representation, viz.:

1. For a given face *A*_*w*_*B*_*w*_*C*_*w*_ in the white surface, and its corresponding face *A*_*p*_*B*_*p*_*C*_*p*_ in the pial surface, define an oblique truncated triangular pyramid.
2. Split this truncated pyramid into three tetrahedra, defined as:

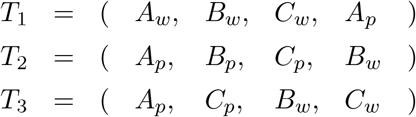 This division leaves no volume under- or over-represented.
3. For each such tetrahedron, let **a**, **b**, **c** and **d** represent its four vertices in terms of coordinates [*x y z*]′. Compute the volume as |**u** · (**v** × **w**)|/6, where **u** = **a** − **d**, **v** = **b** − **d**, **w** = **c** − **d**, the symbol × represents the cross product, · represents the dot product, and the bars | | represent the vector norm.

Computation can be accelerated by setting **d** = *A*_*p*_, the common vertex for the three tetrahedra, such that the vector subtractions can happen only once. Conversion from facewise volume to vertexwise is possible, and done in the same manner as for facewise area. The above method for measuring volume has become the default in the current FreeSurfer version (6.0.0).

### 2.3. Spherical transformation and registration

The white surface is homeomorphically transformed to a sphere (Fischl et al., 1999b), thus keeping a one-to-one mapping between vertices of the native geometry and the sphere. Various strategies are available to place these surfaces in register with a common reference and allow inter-subject comparisons, including the method used by FreeSurfer (Fischl et al., 1999b), Spherical Demons (SD) (Yeo et al., 2010), Multimodal Surface Matching (MSM) (Robinson et al., 2014), among others. Methods that are not diffeomorphic by design but in practice produce invertible and smooth warps can, in principle, be used in registration for areal analyses. In the present analyses, FreeSurfer was used (a complementary comparison with SD is shown in Supplementary Material §2). The measurements of interest obtained from native geometry or in native space, such as area and thickness, are stored separately and are not affected by the spherical transformation or registration.

### 2.4. Interpolation methods

Statistical comparisons require meshes with a common resolution where each point represents homologous locations across individuals. A geodesic sphere has many advantages for this purpose: ease of computation, edges of roughly similar sizes and, if the resolution is fine enough, edge lengths that are much smaller than the diameter of the sphere (Kenner, 1976). We compared four different interpolation methods, described below, at each of three different mesh resolutions: IC3 (642 vertices and 1280 faces), IC5 (10242 vertices and 20480 faces), and IC7 (163842 vertices and 327680 faces); for the comparison between VLBW and controls, the resolution used was IC7, with nearest neighbour interpolation.

#### Nearest neighbour

The well-known nearest neighbour interpolation does not guarantee preservation of areal quantities, although modifications can be introduced to render it approximately mass conservative: for each vertex in the target, the closest vertex is found in the source sphere, and the area from the source vertex is assigned to the target vertex; if a given source vertex maps to multiple target vertices, its area is divided between them so as to preserve the total area. If there are any source vertices that have not been represented in the target, for each one of these, the closest target vertex is located and the corresponding area from the source surface is incremented to any area already stored on it. This method ensures that total area remains unchanged after mapping onto the group surface. This process is a surface equivalent of Jacobian correction^2^ used in volume-based methods in that it accounts for stretches and shrinkages while preserving the overall amount of areal quantities. Nearest neighbour interpolation is the default method in FreeSurfer.

#### Retessellation of the native geometry

This method appeared in Saad et al. (2004). It consists of generating a new mesh by interpolating the coordinates of the vertices in the native geometry to the common grid, thus defining a new surface. The coordinates of each vertex can be treated as a single vector and barycentric interpolation performed in a single step, as:

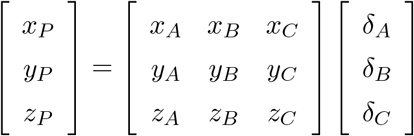
where *x*, *y*, *z* represent the coordinates of the triangular face *ABC* and of the interpolated point *P*, all in native geometry, and *δ* are the barycentric coordinates of *P* with respect to the same face after the spherical transformation. Among the four methods considered in this chapter, this is the only one that does not directly interpolate either area or areal quantities, but the mesh. The area for each face or vertex can then be computed from the new mesh and used for statistical analyses.

#### Redistribution of areas

This method works by splitting the areal quantity present at each vertex in the source sphere using the proportion given by the barycentric coordinates of that vertex in relation to the face on which it lies in the target sphere, thus redistributing these quantities to the three vertices that constitute that face in the target. If some quantity was already present in the target vertex (e.g., from other source vertices lying on the same target face), that quantity is incremented. This method can be represented as:

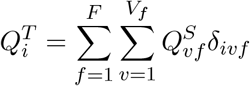
where 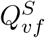 is the areal quantity in the source vertex *v*, *v* ∈ {1,…,*V*_*f*_} lying on the target face *f*, *f* ∈ {1,…,*F*}, *F* being the number of faces that meet at the target vertex *i*, and *δ*_*ivf*_ is the barycentric coordinate of *v*, lying on face *f*, and in relation to the target vertex *i*. The key difference between this method and the classical barycentric interpolation is that, in the latter, the coordinates of the target vertex in relation to their containing source face are used to weight the quantities, in a process that is not mass conservative. Here barycentric coordinates of the source vertex in relation to their containing target face are used; the quantities are split proportionately, and redistributed across target vertices.

#### Pycnophylactic

The ideal interpolation method should conserve the areal quantities globally, regionally and locally, that is, the method has to be *pycnophylactic*. This is accomplished by assigning, to each face in the target sphere, the areal quantity of all overlapping faces from the source sphere, weighted by the fraction of overlap between them (Markoff and Shapiro, 1973; Winkler et al., 2012). The pycnophylactic method operates directly on the faces, not on vertices, and the area (or any other areal quantity) is transferred from source to target surface via weighting by the overlapping area between any pairs of faces. The interpolated areal quantity, 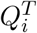, of a face *i* in the target surface, that overlaps with *F* faces from the source surface, is given by:

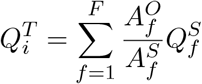
where 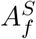 is the area of the *f*-th overlapping face from the source sphere, which contains a quantity 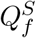 of some areal measurement (such as the surface area measured in the native space), and 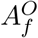 is the overlapping area with the face *i*.

### 2.5. Smoothing

For the comparison of the areal interpolation methods and for the volume methods, smoothing was applied at two levels: no smoothing, and smoothing with a Gaussian kernel with full width at half maximum (FWHM) of 10 mm, chosen so as to preserve the effect of the different resolutions being investigated. For the comparison between VLBW and controls, 30 mm, as in Rimol et al. (2016). Before smoothing, correction for unequal face sizes (Winkler et al., 2012) was applied for all interpolation methods.

### 2.6. Statistical analysis

The statistical analysis between VLBW and controls was performed using PALM - Permutation Analysis of Linear Models (Winkler et al., 2014). The number of permutations was set to 1000, followed by approximation of the tail of the distribution by a generalised Pareto distribution (GPD; Winkler et al., 2016a). Familywise error rate correction (FWER) was done considering both hemispheres and both test directions for the null hypothesis of no difference between the two groups. Analyses were performed separately for cortical thickness, area, and volume (both methods), and also using NPC with Fisher’s combination of p-values (Fisher, 1932) for the joint analysis of thickness and area; Supplementary Material §5 shows an overview of how these analyses are related, whereas details of NPC for imaging data are found in Winkler et al. (2016b).

## 3. Results

### 3.1. Preservation of areal quantities

All methods preserved generally well the global amount of surface area, and therefore, of other areal quantities, at the highest resolution of the common grid (IC7). At lower resolutions, massive amounts of area were lost with the retessellation method: about 40% on average for IC3 (lowest resolution, with 642 vertices and 1280 faces) and 9% for IC5 (intermediate resolution, with 10242 vertices and 20480 faces), although only 1% for IC7 (163842 vertices and 327680 faces). Areal losses, when existing, tended to be uniformly distributed across the cortex (Figure 3, upper panels), with no trends affecting particular regions and, except for retessellation, could be substantially alleviated by smoothing. With the latter, areal losses accumulated throughout the cortex, and the global cortical area, if computed after interpolation, became substantially reduced (biased downwards), even at the highest resolution of the common grid. An extended set of results that support these findings is shown in Supplementary Material §2.

**Figure 3:**
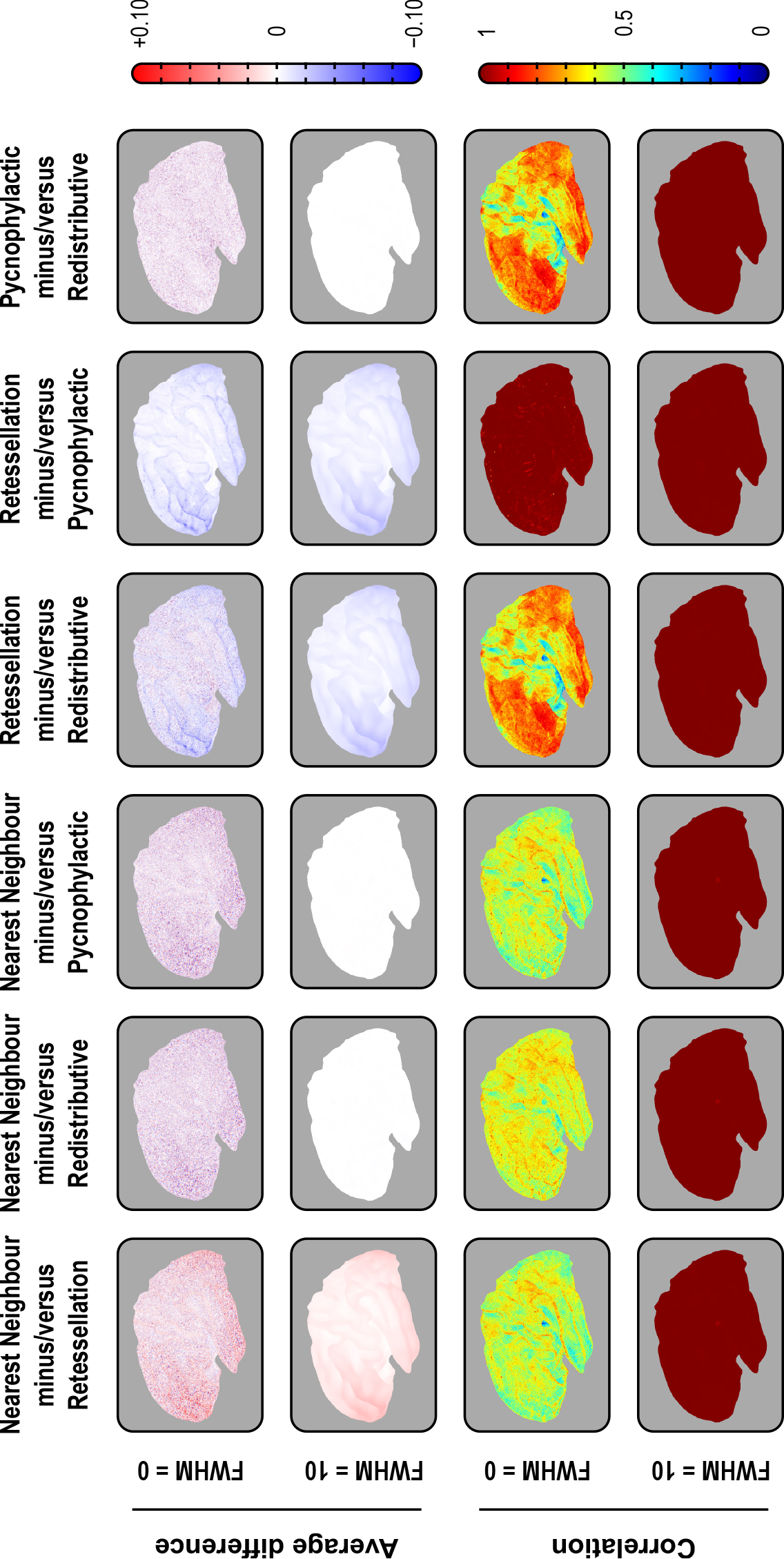
Pairwise average differences (in mm^2^) and correlations between the four interpolation methods, using the IC7 as target, with or without smoothing with a Gaussian kernel of FWHM = 10 mm, projected to the average white surface. Although the four methods differ, with some leading to substantial, undesirable losses and gains in surface area, and the introduction of noise manifested by lower correlations, the average variation was zero for nearest neighbour, redistributive and pycnophylactic. The retessellation method led to substantial losses of area that could not be recovered or compensated by blurring. Although this method showed excellent correlation with pycnophylactic, quantitative results after interpolation are biased downwards. For the medial views, for the right hemisphere, for IC3 and IC5, and for projections to the pial and inflated surfaces, consult the Supplemental Material.

### 3.2. Differences between interpolation methods

While there were no spatial trends in terms of areal gains or losses, the inexactness of the non-pycnophylactic interpolation methods introduced noise that substantially reduced their correlation when assessed between subjects (Figure 3, lower panels). The only exception was between the retessellation and the pycnophylactic methods, which had near perfect correlation even without any smoothing. Smoothing increased the correlation between all methods to near unity throughout the cortex (Supplementary Material §2a). At the subject level, the spatial correlation between the nearest neighbour and the pycnophylactic method was only about 0.60, although approached unity when the subjects were averaged (Supplementary Material §2b). Smoothing lead to a dramatic improvement of agreement between the methods, causing nearest neighbour to be nearly indistinguishable from the pycnophylactic method. The redistributive method performed in a similar manner, although with a higher correlation without smoothing, i.e., about 0.75 (Supplementary Material §2b).

### 3.3. Cortical volume measurements

At the local scale, differences between the product and the analytic methods of volume estimation were as high as 20% in some regions, an amount that could not be alleviated by smoothing or by changes in resolution. As predicted by Figure 1, differences were larger in the crowns of gyri and depths of sulci, in either case with the reverse polarity (Figure 4, upper panels). The vertexwise correlation between the methods across subjects, however, was in general very high, approaching unity throughout the whole cortex, with or without smoothing, and at different resolutions. In regions of higher sulcal variability, however, the correlations were not as high, sometimes as low as 0.80, such as in the insular cortex and at the confluence of parietooccipital and calcarine sulci, between the lingual and the isthmus of the cingulate gyrus (Figure 4, lower panels). At least in the case of the insula, this effect may be partly attributed to a misplacement of the white surface in the region lateral to the claustrum (Glasser et al., 2016). Supplementary Material §3 includes additional results that support these findings.

**Figure 4:**
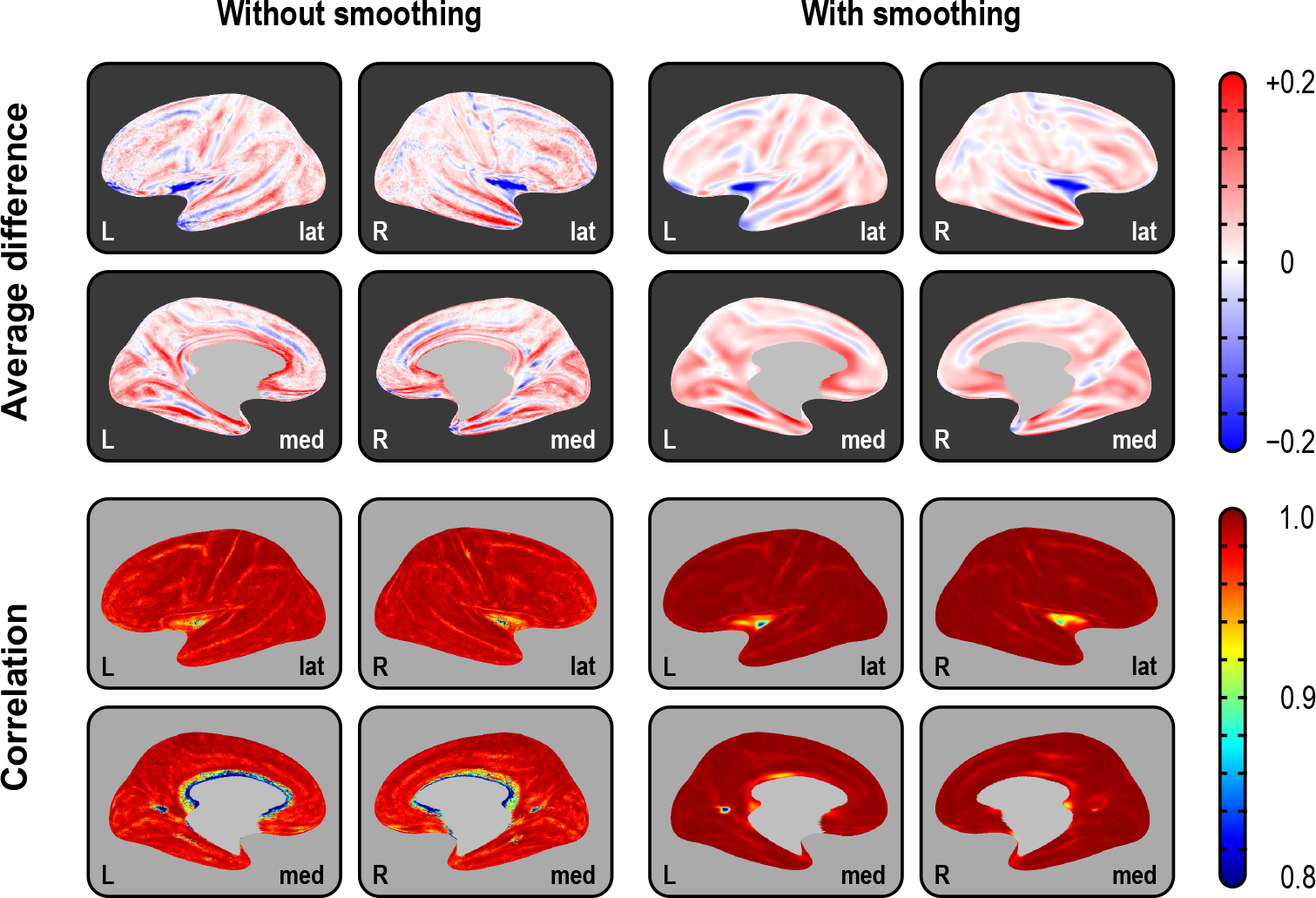
Average difference (in mm^3^) between the two methods of assessing volume and their correlation (across subjects), using the highest resolution (IC7) as the interpolation target, projected to the average inflated surface. As predicated from Figure 1, differences are larger in the crowns of gyri and in the depths of sulci, with gains/losses in volume in these locations following opposite patterns. Although the correlations tend to be generally high, and increase with smoothing, they are lower in regions of higher inter-individual morphological variability, such as at the anterior end of the cuneus, and in the insular cortex. For IC3 and IC5, and for projections to the white and pial surfaces, consult the Supplemental Material.

### 3.4. Global measurements and their variability

Average global cortical area, thickness, and volume (using both methods) across subjects in the sample are shown in Table 2. Cortical volumes assessed with the multiplicative method were significantly higher (*p* < 0.0001) than using the analytic method. Variability for area was higher than for thickness, and even higher for volume: the average coefficient of variation across subjects (100 · *σ/μ*) was, respectively, 9.9%, 3.2% and 10.5%, after adjusting for group, age, and sex, with the parietal region (bilateral) being the most variable for all measurements. The corresponding spatial maps, as well as correlation and Bland-Altman plots, are shown in Supplementary Material §4.

**Table 2:** Average ± standard deviation of area (in mm^2^), thickness (in mm) and volume (in mm^3^) for the subjects of the control group. Volumes are shown assessed using the multiplicative (*m*) and analytic (*a*) methods; their difference is also shown.

| Measure | Left hemisphere | Right hemisphere | Both hemispheres |
| --- | --- | --- | --- |
| Area | $100877.7 \pm 7868.3$ | $101725.3 \pm 8101.4$ | $202603.0 \pm 15944.4$ |
| Thickness | $2.5556 \pm 0.0896$ | $2.5436 \pm 0.0880$ | $2.5495 \pm 0.0876$ |
| Volume<sup>($m$)</sup> | $257781.8 \pm 21973.5$ | $258751.1 \pm 22694.0$ | $516533.0 \pm 44562.1$ |
| Volume<sup>($a$)</sup> | $254053.8 \pm 21740.6$ | $255181.4 \pm 22441.4$ | $509235.1 \pm 44080.0$ |
| Difference<sup>($m-a$)</sup> | $3728.0 \pm 581.6$ | $3569.8 \pm 624.1$ | $7297.8 \pm 1108.1$ |

### 3.5. Differences between VLBW and controls

Analysing cortical thickness and area separately, the comparisons between VLBW subjects and term-born controls suggested a distinct pattern of differences. Surface area maps showed a significant bilateral reduction in the middle temporal gyrus, the superior banks of the lateral sulcus, and the occipito-temporal lateral (fusiform) gyrus, as well as a diffuse bilateral pattern of areal losses affecting the superior frontal gyrus, posterior parietal cortex and, in the right hemisphere, the subgenual area of the cingulate cortex. Cortical thickness maps showed a diffuse bilateral thinning in the parietal lobes, left middle temporal gyrus, right superior temporal sulcus, while showing bilateral thickening of the medial orbito-frontal cortex and the right medial occipital cortex of the VLBW subjects compared to controls (Figure 5, upper panels, light blue background). Maps of cortical volume differences largely mimicked the surface area results, albeit with a few differences: diffuse signs of volume reduction in the parietal lobes, ascribable to cortical thinning and, contrary to the analysis of area and thickness, no effects found in the medial-orbitofrontal or in the subgenual region of the cingulate gyrus (Figure 5, middle panels, light red background).

**Figure 5:**
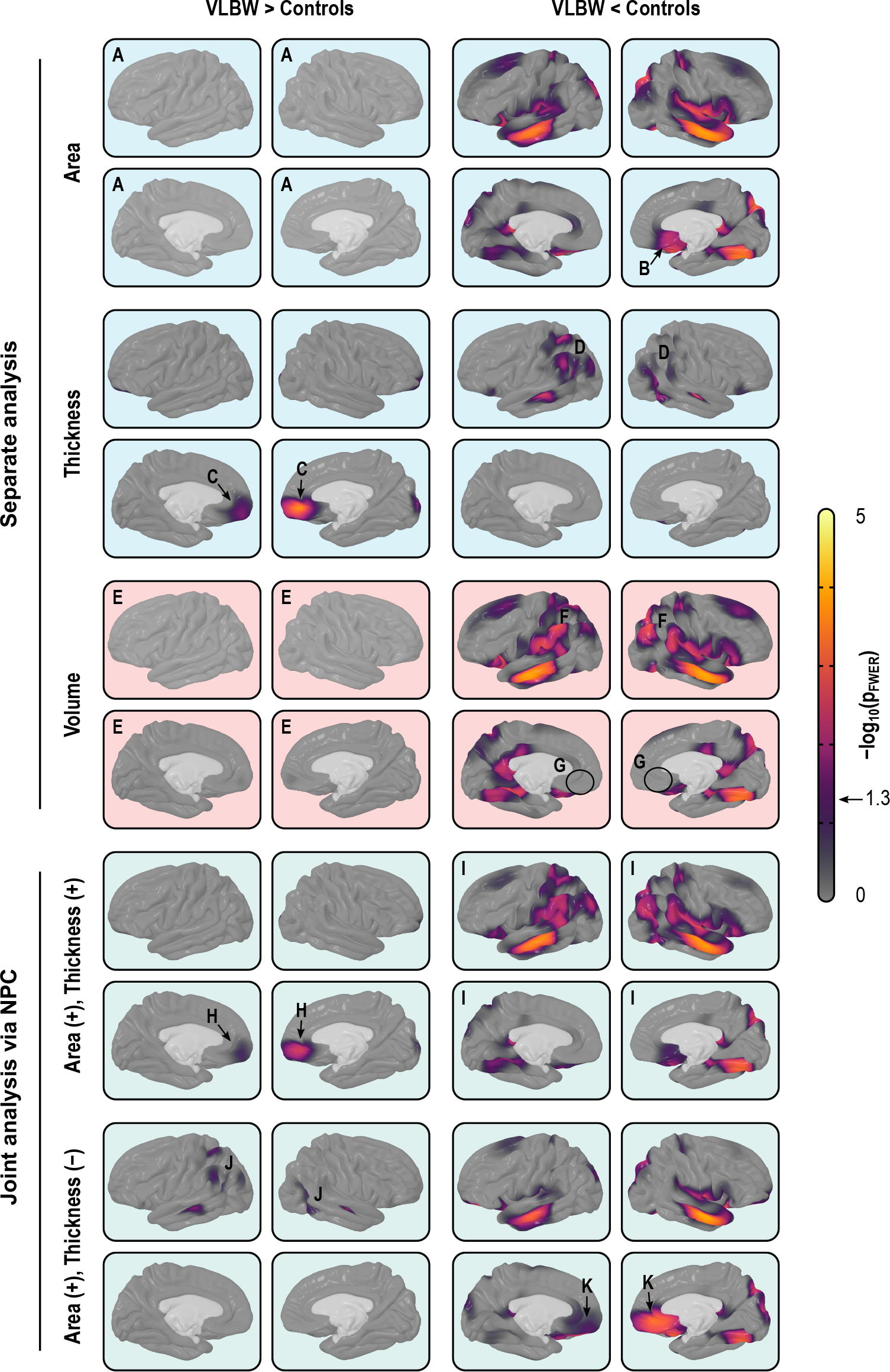
Separate (*light blue background*) and joint (*green*) analysis of cortical area and thickness, as well as volume (*red*), using the IC7 resolution and smoothing with FWHM = 30mm. Analysis of area indicates no increases in the VBLW group anywhere in the cortex (A), and reductions in, among other regions, the subgenual region of the cingulate cortex (B). Analysis of thickness indicates that VBLW subjects have thicker cortex in the medial orbitofrontal cortex (C) and in the right medial occipital cortex, as well as diffuse bilateral thinning in parietal and middle temporal regions (D). Analysis of volume alone broadly mimics analysis of area, with no evidence of increased volume in vlbw subjects (E), although in some maps there seems to be a partial superimposition of the effects seen separately for area and thickness, with signs of bilaterally decreased volume throughout the parietal lobe (F), but contrary to the analysis of area, no signs of reduction in the subgenual cortex (G). Jointly analysing area and thickness gives equal weight to both measurements, and allows directional effects to be inferred. Contrary to the case for volume, it is possible to know that there is an increase in the amount of cortical tissue in vlbw subjects in the medial orbito-frontal cortex (H) when compared to controls, and a bilateral decrease throughout most of the parietal cortex, stronger in the middle temporal and fusiform gyri, in both hemispheres (I). Moreover, the joint analysis allows search for effects that can negate each other, such as in this case weaker effects in the parietal region (J), that partially overlap in space with those shown in (I). Finally, strong effects in the middle orbitofrontal, that were missed with simple volumes (G) become clearly visible (K).

### 3.6. Joint analysis via NPC

Non-parametric combination of thickness and area provided information about patterns of group differences not visible in cortical volume analyses (Figure 5, lower panels, light green background). In the present data, the joint analysis suggested a decrease in the amount of tissue in VLBW subjects in the medial orbito-frontal cortex, which was not visible in the volume analysis, as well as a bilateral decrease throughout most of the parietal cortex, and in the middle temporal and fusiform gyri. Finally, NPC showed simultaneous bilateral decrease in surface area and increase in thickness in the medial orbito-frontal gyrus, none of which was observed using simple volume measurements. Additional maps are shown in Supplementary Material §5.

## 4. Discussion

### 4.1. Interpolation of areal quantities

The different interpolation methods did not perform similarly in all settings. Nearest neighbour and redistributive required smoothing of at least FWHM = 10 mm as used here in order to become comparable to, and interchangeable with, the pycnophylactic method. However, since data is usually smoothed in neuroimaging studies in order to improve the matching of homologies and to improve the signal-to-noise ratio, this is not a significant limitation. Retessellation, particularly at lower resolutions, lead to substantial areal losses that could not be recovered even with smoothing. Moreover, the vertices of the retessellated surfaces are not guaranteed to lie at the tissue boundaries they represent, introducing uncertainties to the obtained measurements. Regarding speed, although the various implementations run in linear time, the pycnophylactic method has to perform a larger number of computations that may not pay off when compared with nearest neighbour, provided that smoothing is used.

### 4.2. Volumes improved, yet problematic

The large absolute difference between the product and the analytic method for cortical volume indicates that if interest lies in the actual values (for instance, for predictive models), the analytic method is to be preferred. The high correlation across subjects, however, suggests that, for group comparisons and similar analyses, both methods generally lead to similar results, except in a few regions of higher morphological inter-individual variability. However, even for group comparisons, cortical volume is a poor choice of trait of interest. Even though volume encapsulates information from both area and thickness, research has suggested that the proportion in which the variability of these two measurements coalesces varies spatially across the cortical mantle (Winkler et al., 2010; Storsve et al., 2014). Moreover, previous literature suggests that most of the between-subject variability in cortical volume, including that measured using VBM, can be explained by the variability of surface area (Voets et al., 2008; Lenroot et al., 2009; Winkler et al., 2010; Rimol et al., 2012), whereas most of the within-subject variability can be explained by variability of cortical thickness, at least during adult life (Storsve et al., 2014), thus rendering volume a largely redundant metric. In effect, the continuous cortical maps in Figure 5, resulting from a between-subject analysis, confirm that the results for cortical volume largely mirror the results for cortical surface area.

### 4.3. Joint analyses via NPC

Such problems with cortical volume can be eschewed through the use of a joint statistical analysis of area and thickness. The NPC methodology gives equal (or otherwise predefined) weights for thickness and area, which therefore no longer have their variability mixed in unknown and variable proportions across the cortical mantle. Various combining functions can be considered, and the well-known Fisher method of combination of p-values is a simple and computationally efficient choice. By using two distinct metrics in a single test, power is increased (Fisher, 1932; Pesarin and Salmaso, 2010b; Winkler et al., 2016b), allowing detection of effects that otherwise may remain unseen when analysing volume, or when thickness and area are used separately. NPC can be particularly useful for the investigation of processes affecting cortical area and thickness simultaneously, even if in opposite directions or at different rates (both phenomena that have been recently reported, e.g., Hogstrom et al., 2013; Storsve et al., 2014), and can effectively replace volume as the measurement of interest in these cases, with various benefits and essentially none of the shortcomings. It constitutes a general method that can be applied to any number of partial tests, each relating to hypotheses on data that may be of a different nature, obtained using different measurement units, and related to each other arbitrarily.

Moreover, NPC allows testing directional hypotheses (by reversing the signs of partial tests), hypotheses with concordant directions (taking the extremum of both after multiple testing correction), and two-tailed hypotheses (with two-tailed partial tests). Power increases consistently with the introduction of more partial tests when there is a true effect, while strictly controlling the error rate. This is in contrast to classical multivariate tests based on regression, such as MANOVA or MANCOVA, that do not provide information on directionality of the effects, and lose power as the number of partial tests increase past a certain optimal point. Usage of NPC is not constrained to the replacement of cortical volume, and the method can be considered for analyses involving other cortical indices, including myelination (Glasser and Van Essen, 2011; Sereno et al., 2013) and folding and gyrification metrics (Mangin et al., 2004; Schaer et al., 2008; Toro et al., 2008) that can interact in distinct and complex ways (Tallinen et al., 2014, 2016), among others. A joint analysis offers an elegant solution to the problem of multiple comparisons, both across locations on the cortical surface (vertices), due to its non-parametric nature, and across measurements; it also offers increased power over separate analyses.

### 4.4. Permutation inference

NPC is based on permutations in each of the partial tests but does not preclude separate analysis of thickness and area, and can accommodate partial tests combining positive and negative directions. Using permutation tests with synchronized shuffling, it is trivial to correct for the multiplicity of tests while taking their non-independence into account. Permutation tests provide exact inference based on minimal assumptions, while allowing multiple testing correction with strong control over the error rate. Even though these tests still have certain requirements, such as the data being exchangeable, various types of structured dependency can be accommodated by means of restricted permutation strategies. Finally, permutation tests do not depend on distributional assumptions, which is an advantage when analysing surface area, since area at the local level shows positive skewness and is better characterised as log-normal (Winkler et al., 2012).

### 4.5. Area and thickness of VLBW subjects

The sample used for this analysis is particularly suitable as neurodevelopmental brain disorders associated with preterm birth are known to have a divergent effect on cortical area and cortical thickness, including both cortical thinning and thickening (Rimol et al., 2016), hence a joint analysis being potentially more informative in lieu of simple cortical volume. Here, the reduced cortical surface area observed in VLBW subjects compared to controls replicates previous findings from the same cohort at 20 years of age (Skranes et al., 2013), and is consistent with findings from a younger cohort of VLBW subjects (Sølsnes et al., 2015) and teenagers born with extremely low birth weight (⩽ 1000g) (Grunewaldt et al., 2014). The combined evidence from these studies suggests that surface area reductions in the preterm brain are present from early childhood and remain until adulthood (Rimol et al., 2016), and various mechanisms have been proposed (Volpe, 2009, 2011; Eikenes et al., 2011; Hagberg et al., 2015). Likewise, explanations for cortical thinning in some regions of VLBW subjects, but the thickening of others have been proposed (Marín-Padilla, 1997; Bjuland et al., 2013; Grunewaldt et al., 2014). The combination of thickening and reduced area in medial orbito-frontal cortex has been observed in multiple cohorts and, on the light of previously proposed mechanisms, these changes could be related to prenatal factors, such as foetal growth restriction, or to postnatal exposure to extra-uterine environmental stressors (Sølsnes et al., 2015; Rimol et al., 2016). Regardless of underlying pathological aspects, the morphological indices appear to be robust markers of perinatal brain injury and maldevelopment (Raznahan et al., 2011; Skranes et al., 2013; Rimol et al., 2016).

### 4.6. Limitations

As NPC is a permutation test, the assumption of exchangeability must hold, which can be a limitation when certain types dependencies between observations exist. The method can be computationally intensive, particularly for datasets that are large or have high resolution. Both problems can be addressed, at least in particular cases: structured dependencies (such as when studying twins) can be accommodated by imposing restrictions on which permutations can be performed (Winkler et al., 2015), whereas accelerations can be accomplished using various approximate or exact methods (Winkler et al., 2016a); the latter were used in this particular analysis. Regarding power, the present VLBW sample is medium-sized, and it is possible that real group effects could not be detected, including volume differences. However, to the extent that cortical thickness and surface area go in opposite directions, failure to detect group differences in cortical volume can be unrelated to statistical volume power issues.

## 5. Conclusion

We studied the four extant interpolation methods for the assessment of cortical area, and observed that the nearest neighbour interpolation, followed by a Jacobian correction and smoothing, is virtually indistinguishable from the pycnophylactic method, while offering reduced computational costs. This leads us to recommend, for practical purposes, the nearest neighbour method, with smoothing, when investigating cortical surface area. In addition, we demonstrated that the non-parametric combination of cortical thickness and area can be more informative than a simple analysis of cortical volume, even when the latter is assessed using the improved, analytic method that does not over or under-represent tissue according to the cortical convolutions.

## Supplementary Material

Detailed results have been organised in a set of browsable pages and packaged into a 7.8 GB file that constitutes the Supplementary Material. It is deposited for long term preservation and public access at the Dryad Digital Repository, under the Digital Object Identifier (DOI): 10.5061/dryad.hlv85. The results above make ample reference to this material, and its inspection is encouraged. The Supplementary Material also includes high resolution and complementary views of all figures shown in the main text. A mirrored copy that does not require download, though not guaranteed for permanent preservation, can be found at http://bit.ly/2x9F96b.

## Acknowledgements

The authors thank Donald J. Hagler Jr. (University of California, San Diego, CA, USA), and David Van Essen and Matthew F. Glasser (Washington University Medical School, St. Louis, MO, USA) for the helpful discussions at different stages of this work. This research project was supported by grants from the Research Council of Norway (ES182663) and the Central Norway Regional Health Authority (46056610 and 46039500). A.M.W. received supported by the National Research Council of Brazil (CNPq; 211534/2013-7). T.E.N. is supported by the Wellcome Trust (100309/Z/12/Z). M.R.S. received funding from the NIH R21AG050122-01A1, R41AG052246-01 and 1K25EB013649-01 projects. Data processing was largely performed on the Abel Cluster, owned by the University of Oslo and the Norwegian Metacenter for High Performance Computing (NOTUR), and operated by the Department for Research Computing at USIT, University of Oslo.

## Footnotes

1 Available at freesurfer.net

2 Not to be confused with the computation of the Jacobian itself, that is defined, for the *i*-th vertex, as 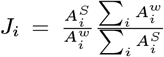, where 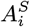 is the area of the vertex in the source (registered) sphere, 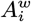 is the area of the same vertex in the white surface (native space and native geometry), and the sums are over the entire surface, i.e., all vertices.

